# Identification of new therapeutic targets for osteoarthritis through genome-wide analyses of UK Biobank

**DOI:** 10.1101/453530

**Authors:** Ioanna Tachmazidou, Konstantinos Hatzikotoulas, Lorraine Southam, Jorge Esparza-Gordillo, Valeriia Haberland, Jie Zheng, Toby Johnson, Mine Koprulu, Eleni Zengini, Julia Steinberg, Jeremy M Wilkinson, Sahir Bhatnagar, Joshua Hoffman, Natalie Buchan, Dániel Süveges, arcOGEN Consortium, Laura Yerges Armstrong, George Davey Smith, Tom R Gaunt, Robert A Scott, Linda C McCarthy, Eleftheria Zeggini

## Abstract

Osteoarthritis is the most common musculoskeletal disease and the leading cause of disability globally. Here, we perform the largest genome-wide association study for osteoarthritis to date (77,052 cases and 378,169 controls), analysing 4 phenotypes: knee osteoarthritis, hip osteoarthritis, knee and/or hip osteoarthritis, and any osteoarthritis. We discover 64 signals, 52 of them novel, more than doubling the number of established disease loci. Six signals fine map to a single variant. We identify putative effector genes by integrating eQTL colocalization, fine-mapping, human rare disease, animal model, and osteoarthritis tissue expression data. We find enrichment for genes underlying monogenic forms of bone development diseases, and for the collagen formation and extracellular matrix organisation biological pathways. Ten of the likely effector genes, including *TGFB1*, *FGF18*, *CTSK* and *IL11* have therapeutics approved or in clinical trials, with mechanisms of action supportive of evaluation for efficacy in osteoarthritis.

---

Osteoarthritis affects 40% of individuals over the age of 70^1^, is a major cause of pain, comorbidity and mortality^2^. Ten million people in the UK alone suffer from osteoarthritis, with a total indirect cost to the economy of £14.8 billion per annum^2^. Disease management targets the main symptom (pain) and culminates in joint replacement surgery (1.76 million per year in the EU) with variable outcomes^3^. There is a clear and urgent need to translate genomic evidence into druggable mechanisms of disease aetiology and progression, to support the development of disease-modifying therapies for osteoarthritis.

Here, we leverage the UK Biobank and arcOGEN resources to perform the largest genome-wide meta-analysis for osteoarthritis to date across ∼17.5 million single nucleotide variants in up to 455,221 individuals (Supplementary Figure 1). We identify 65 genome-wide significant variants at 64 loci (*P*≤3×10^-8^; Online Methods, Supplementary Table 1), 52 of which are novel, thus increasing the number of established loci from 34^4^ to 86: 24 novel signals for osteoarthritis at any site (77,052 cases), 15 for hip osteoarthritis (15,704 cases), 7 for knee osteoarthritis (24,955 cases), and 6 for osteoarthritis of the hip and/or knee (39,427 cases) (Table 1; Supplementary Table 2; Supplementary Figures 2-6). We find that 25 of 34 previously-reported loci show association (*P*<0.05) with at least one of the four osteoarthritis traits we evaluate (Supplementary Table 3).

**Table 1:** **Independent variants with *P* <3×10 ^-8^ in an inverse-variance weighted fixed effects meta-analysis of UK Biobank and arcOGEN.** Variant positions are reported according to build 37 and their alleles are coded based on the positive strand.

| SNPID | RSID | Most Severe Consequence | Trait | Other Traits GWAS sign meta-analysis | EA | NE A | UK Biobank |  | arcOGEN |  | WEAF | OR | OR_L 95 | OR_U95 | PV | q_pV | i2 | EAF |  |
| --- | --- | --- | --- | --- | --- | --- | --- | --- | --- | --- | --- | --- | --- | --- | --- | --- | --- | --- | --- |
|  |  |  |  |  |  |  | OR | PV | OR | PV |  |  |  |  |  |  |  | Cases/Controls * | 1000 Genomes GBR ** |
| Newly Identified Loci |  |  |  |  |  |  |  |  |  |  |  |  |  |  |  |  |  |  |  |
| 1:103572927 | rs4338381 | sequence_feature | OA_hip | OA;OA_kneehip | A | G | 1.1 | 4.70E-13 | 1.1 | 2.71E-03 | 0.63 | 1.1 | 1.07 | 1.13 | 4.37E-15 | 0.93 | 0 | 0.6529/0.6307<br>0.654/0.6316 | 0.62<br>(0.55, 0.69) |
| 1:150214028 | 1:150214028 | downstream_gene | OA | OA_hip | C | CT | 1.03 | 2.50E-08 |  |  | 0.37 | 1.03 | 1.02 | 1.05 | 2.54E-08 | 1 | 0 | 0.3679/0.36<br>- | - |
| 1:174192402 | 1:174192402 | intron | OA |  | TAAAAA<br>AAAAA<br>AAAAA<br>AA | T | 1.03 | 1.10E-08 |  |  | 0.57 | 1.03 | 1.02 | 1.05 | 1.05E-08 | 1 | 0 | 0.5855/0.5758<br>- | - |
| 1:183906245 | rs11583641 | 3_prime_UTR | OA_hip |  | C | T | 1.09 | 3.10E-09 | 1.07 | 5.55E-02 | 0.72 | 1.08 | 1.06 | 1.11 | 5.58E-10 | 0.63 | 0 | 0.7404/0.7246<br>0.7347/0.7218 | 0.68<br>(0.61, 0.74) |
| 1:245750932 | rs10218792 | intron | OA |  | G | T | 1.04 | 7.40E-08 | 1.04 | 5.13E-02 | 0.27 | 1.04 | 1.02 | 1.05 | 2.03E-08 | 0.77 | 0 | 0.2739/0.2675<br>0.274/0.2656 | 0.2<br>(0.15, 0.27) |
| 2:33434336 | rs2061027 | intron | OA | OA_knee;<br>OA_kneehip | A | G | 1.04 | 9.70E-12 | 1.07 | 6.58E-03 | 0.51 | 1.04 | 1.03 | 1.05 | 3.16E-13 | 0.25 | 25.8 | 0.5141/0.5041<br>0.5178/0.5011 | 0.55<br>(0.47, 0.62) |
| 2:192671981 | rs12470967 | intron | OA_knee | OA_kneehip | A | G | 1.06 | 1.50E-08 |  |  | 0.43 | 1.06 | 1.04 | 1.08 | 1.50E-08 | 1 | 0 | 0.4416/0.4291<br>- | 0.55<br>(0.48, 0.63) |
| 2:204387482 | rs62182810 | intron | OA |  | A | G | 1.03 | 6.20E-09 | 1.04 | 5.44E-02 | 0.55 | 1.03 | 1.02 | 1.05 | 1.65E-09 | 0.84 | 0 | 0.554/0.5462<br>0.5562/0.5468 | 0.52<br>(0.45, 0.6) |
| 3:50022049 | rs62262139 | intron | OA |  | A | G | 1.04 | 9.10E-11 |  |  | 0.54 | 1.04 | 1.03 | 1.05 | 9.09E-11 | 1 | 0 | 0.544/0.5348<br>- | 0.54<br>(0.47, 0.62) |
| 4:1704244 | rs11732213 | intron | OA_kneehip | OA;OA_hip | T | C | 1.06 | 2.30E-09 | 1.04 | 1.46E-01 | 0.81 | 1.06 | 1.04 | 1.08 | 8.81E-10 | 0.6 | 0 | 0.8142/0.8047<br>0.8159/0.8092 | 0.73<br>(0.66, 0.79) |
| 4:13039440 | rs1913707 | intergenic_region | OA_hip | OA | A | G | 1.07 | 9.00E-08 | 1.16 | 4.24E-06 | 0.61 | 1.08 | 1.06 | 1.11 | 2.96E-11 | 0.03 | 79 | 0.6271/0.6111<br>0.6421/0.6083 | 0.59<br>(0.52, 0.66) |
| 4:25408838 | rs34811474 | missense | OA |  | G | A | 1.04 | 3.70E-09 | 1.03 | 1.93E-01 | 0.77 | 1.04 | 1.03 | 1.05 | 2.17E-09 | 0.76 | 0 | 0.7745/0.7671<br>0.7873/0.7821 | 0.71<br>(0.64, 0.77) |
| 4:103188709 | rs13107325 | missense | OA |  | T | C | 1.1 | 4.10E-18 | 1.08 | 5.14E-02 | 0.08 | 1.1 | 1.07 | 1.12 | 8.29E-19 | 0.7 | 0 | 0.0801/0.0735<br>0.0871/0.0811 | 0.1<br>(0.06, 0.15) |
| 5:77467824 | rs35611929 | intron | OA_knee |  | A | G | 1.06 | 5.50E-08 | 1.06 | 7.93E-02 | 0.34 | 1.06 | 1.04 | 1.08 | 1.21E-08 | 0.96 | 0 | 0.3544/0.3411<br>0.3418/0.3296 | 0.32<br>(0.25, 0.39) |
| 5:170871074 | rs3884606 | intron | OA_kneehip |  | G | A | 1.04 | 2.10E-07 | 1.06 | 3.43E-03 | 0.49 | 1.04 | 1.03 | 1.06 | 8.25E-09 | 0.42 | 0 | 0.498/0.4868<br>0.4997/0.4843 | 0.47<br>(0.39, 0.54) |
| 6:26123502 | rs115740542 | upstream_gene | OA | OA_hip;OA_kneehip | C | T | 1.06 | 2.00E-08 | 1.06 | 6.72E-02 | 0.07 | 1.06 | 1.04 | 1.08 | 8.59E-09 | 0.95 | 0 | 0.0776/0.0731<br>0.0743/0.0704 | 0.05<br>(0.02, 0.09) |
| 6:33055501 | rs9277552 | downstream_gene | OA_kneehip | OA_knee;OA | C | T | 1.06 | 9.30E-10 | 1.05 | 1.07E-01 | 0.79 | 1.06 | 1.04 | 1.08 | 2.37E-10 | 0.78 | 0 | 0.7964/0.7862<br>0.8246/0.8171 | 0.7<br>(0.62, 0.76) |
| 6:44449697 | rs12154055 | intergenic_region | OA |  | G | A | 1.03 | 6.90E-09 | 0.99 | 8.95E-01 | 0.61 | 1.03 | 1.02 | 1.04 | 2.71E-08 | 0.1 | 63.5 | 0.6194/0.6117<br>0.6112/0.613 | 0.66<br>(0.59, 0.73) |
| 6:55636940 | rs80287694 | intron | OA_hip |  | G | A | 1.13 | 2.10E-09 | 1.05 | 2.64E-01 | 0.11 | 1.12 | 1.08 | 1.16 | 2.66E-09 | 0.2 | 39.1 | 0.1246/0.1128<br>0.1159/0.1107 | 0.15<br>(0.1, 0.22) |
| 7:95719833 | rs11409738 | intron | OA | OA_kneehip | TA | T | 1.04 | 2.10E-10 |  |  | 0.37 | 1.04 | 1.03 | 1.05 | 2.13E-10 | 1 | 0 | 0.3751/0.3664<br>- | 0.39<br>(0.32, 0.47) |
| 8:9087679 | rs330050 | upstream_gene | OA | OA_kneehip;OA_hip | G | C | 1.04 | 2.90E-10 | 1.06 | 9.51E-01 | 0.51 | 1.04 | 1.03 | 1.05 | 1.93E-11 | 0.35 | 0 | 0.5145/0.5052<br>0.5141/0.4995 | 0.45<br>(0.37, 0.52) |
| 8:130768503 | rs60890741 | upstream_gene | OA_hip |  | C | CA | 1.11 | 4.50E-09 |  |  | 0.86 | 1.11 | 1.08 | 1.16 | 4.50E-09 | 1 | 0 | 0.8796/0.8671<br>- | 0.87<br>(0.81, 0.91) |
| 9:116911147 | rs919642 | intergenic_region | OA | OA_kneehip;OA_knee | T | A | 1.05 | 1.10E-14 | 1.03 | 1.96E-01 | 0.27 | 1.05 | 1.04 | 1.06 | 8.55E-15 | 0.41 | 0 | 0.2697/0.2595<br>0.2774/0.2717 | 0.27<br>(0.21, 0.35) |
| 9:117840742 | rs1330349 | upstream_gene | OA_hip |  | C | G | 1.08 | 2.40E-09 | 1.09 | 3.93E-03 | 0.58 | 1.08 | 1.06 | 1.11 | 4.10E-11 | 0.71 | 0 | 0.5926/0.5738<br>0.5988/0.5773 | 0.6<br>(0.53, 0.68) |
| 9:129386860 | rs62578127 | intron | OA_hip |  | C | T | 1.09 | 2.00E-11 | 1.07 | 2.15E-02 | 0.63 | 1.09 | 1.06 | 1.11 | 2.77E-12 | 0.54 | 0 | 0.6514/0.6307<br>0.6552/0.6401 | 0.6<br>(0.52, 0.67) |
| 11:28874997 | rs17659798 | intron | OA_knee<br>hip |  | A | C | 1.05 | 2.50E-09 | 1.06 | 2.14E-02 | 0.71 | 1.06 | 1.04 | 1.07 | 2.06E-10 | 0.86 | 0 | 0.725/0.7135<br>0.7266/0.7149 | 0.69<br>(0.61, 0.75) |
| 11:30774280 | rs11031191 | interge<br>nic_regi<br>on | OA |  | T | G | 1.03 | 3.30E-08 | 1.03 | 9.65E-02 | 0.35 | 1.03 | 1.02 | 1.05 | 1.42E-08 | 0.95 | 0 | 0.3541/0.347<br>0.3552/0.348 | 0.36<br>(0.29, 0.44) |
| 11:65323725 | rs10896015 | upstream_gene | OA_hip |  | G | A | 1.08 | 2.30E-07 | 1.11 | 2.13E-03 | 0.73 | 1.08 | 1.05 | 1.11 | 2.74E-09 | 0.36 | 0 | 0.7446/0.7299<br>0.7491/0.7283 | 0.69<br>(0.62, 0.76) |
| 11:66501624 | rs34419890 | upstream_gene | OA_hip |  | T | C | 1.13 | 2.70E-07 | 1.16 | 8.15E-03 | 0.93 | 1.13 | 1.09 | 1.18 | 1.99E-08 | 0.75 | 0 | 0.9334/0.9249<br>0.9401/0.9314 | 0.93<br>(0.88, 0.96) |
| 11:76506572 | rs1149620 | upstream_gene | OA |  | T | A | 1.04 | 2.50E-09 | 1.04 | 8.13E-02 | 0.57 | 1.04 | 1.02 | 1.05 | 6.93E-10 | 0.9 | 0 | 0.58/0.5712<br>0.576/0.5668 | 0.58<br>(0.51, 0.65) |
| 12:59289598 | rs79056043 | intron | OA_hip |  | G | A | 1.16 | 9.50E-07 | 1.28 | 6.44E-05 | 0.05 | 1.18 | 1.12 | 1.24 | 1.33E-09 | 0.14 | 53 | 0.0509/0.0444<br>0.0651/0.0516 | 0.06<br>(0.03, 0.11) |
| 12:69637847 | rs317630 | intron | OA |  | T | C | 1.03 | 7.50E-08 | 1.04 | 1.79E-01 | 0.27 | 1.04 | 1.02 | 1.05 | 1.97E-08 | 0.75 | 0 | 0.2786/0.2728<br>0.2841/0.2755 | 0.27<br>(0.21, 0.34) |
| 12:90326919 | rs11105466 | intron | OA_knee<br>hip |  | A | G | 1.04 | 8.30E-07 | 1.07 | 2.37E-03 | 0.42 | 1.04 | 1.03 | 1.06 | 2.15E-08 | 0.26 | 22.<br>6 | 0.4153/0.4152<br>0.4285/0.4118 | 0.41<br>(0.34, 0.48) |
| 12:94167220 | rs2171126 | intron | OA | OA_knee<br>hip | T | C | 1.03 | 1.30E-08 | 1.06 | 6.78E-03 | 0.51 | 1.03 | 1.02 | 1.05 | 9.07E-10 | 0.26 | 21.<br>5 | 0.5137/0.5059<br>0.5193/0.5044 | 0.51<br>(0.44, 0.59) |
| 12:122606837 | rs11059094 | intron | OA_hip |  | T | C | 1.07 | 9.80E-09 | 1.1 | 9.54E-04 | 0.48 | 1.08 | 1.05 | 1.1 | 7.38E-11 | 0.44 | 0 | 0.4943/0.4769<br>0.4967/0.4723 | 0.45<br>(0.37, 0.52) |
| 12:123835233 | rs56116847 | upstream_gene | OA_knee | OA;OA_kn<br>eehip | A | G | 1.07 | 4.40E-11 | 1 | 9.37E-01 | 0.36 | 1.06 | 1.04 | 1.08 | 3.19E-10 | 0.05 | 74.<br>2 | 0.3685/0.3537<br>0.3635/0.3628 | 0.35<br>(0.28, 0.43) |
| 15:50731131 | rs35912128 | upstream_gene | OA_knee |  | AT | A | 1.08 | 2.20E-08 |  |  | 0.17 | 1.08 | 1.05 | 1.11 | 2.18E-08 | 1 | 0 | 0.1717/0.1613<br>- | 0.25<br>(0.19, 0.32) |
| 15:75097780 | rs35206230 | downstream_gene | OA | OA_knee<br>hip | T | C | 1.04 | 8.10E-12 | 1.05 | 6.44E-02 | 0.67 | 1.04 | 1.03 | 1.05 | 1.48E-12 | 0.86 | 0 | 0.6795/0.67<br>0.6725/0.6624 | 0.68<br>(0.61, 0.74) |
| 16:69735271 | rs6499244 | 3_prime_UTR | OA_knee | OA_kneehip | A | T | 1.06 | 6.40E-10 | 1.07 | 5.55E-03 | 0.56 | 1.06 | 1.04 | 1.08 | 3.88E-11 | 0.74 | 0 | 0.4266/0.4414<br>0.5768/0.5591 | 0.52<br>(0.45, 0.6) |
| 16:89704365 | rs1126464 | missense | OA |  | G | C | 1.04 | 3.00E-11 | 0.99 | 8.88E-01 | 0.76 | 1.04 | 1.03 | 1.06 | 1.56E-10 | 0.07 | 69.3 | 0.7638/0.7566<br>0.763/0.7644 | 0.73<br>(0.66, 0.8) |
| 17:2058349 | rs35087650 | intron | OA_knee |  | ATT | A | 1.07 | 1.20E-09 |  |  | 0.26 | 1.07 | 1.05 | 1.1 | 1.18E-09 | 1 | 0 | 0.2548/0.241<br>- | 0.27<br>(0.21, 0.35) |
| 17:29496343 | rs2953013 | intron | OA_kneehip |  | C | A | 1.05 | 1.90E-09 | 1.05 | 4.36E-02 | 0.3 | 1.05 | 1.04 | 1.07 | 3.07E-10 | 0.87 | 0 | 0.3047/0.2138<br>0.3065/0.2964 | 0.27<br>(0.21, 0.34) |
| 17:44038785 | rs62063281 | intron | OA_hip |  | G | A | 1.1 | 1.60E-10 | 1.1 | 9.13E-03 | 0.22 | 1.1 | 1.07 | 1.13 | 5.30E-12 | 0.91 | 0 | 0.2361/0.2198<br>0.2491/0.2321 | 0.25<br>(0.19, 0.32) |
| 17:44057886 | rs547116051 | upstream_gene | OA |  | AC | A | 1.83 | 1.50E-08 |  |  | 0.0013 | 1.83 | 1.49 | 2.26 | 1.50E-08 | 1 | 0 | 0.0006/0.0004<br>- | 0 |
| 17:59652282 | rs7222178 | intergenic_region | OA_hip |  | A | T | 1.1 | 2.60E-10 | 1.08 | 1.23E-02 | 0.2 | 1.1 | 1.07 | 1.13 | 3.78E-11 | 0.59 | 0 | 0.2129/0.197<br>0.2049/0.1926 | 0.18<br>(0.13, 0.25) |
| 17:70012939 | rs8067763 | intergenic_region | OA_knee |  | G | A | 1.06 | 2.50E-09 | 1.03 | 3.83E-01 | 0.41 | 1.06 | 1.04 | 1.08 | 2.39E-09 | 0.35 | 0 | 0.4179/0.4041<br>0.4126/0.4055 | 0.44<br>(0.37, 0.51) |
| 18:20970706 | rs10502437 | intron | OA |  | G | A | 1.03 | 3.60E-08 | 1.02 | 3.39E-01 | 0.6 | 1.03 | 1.02 | 1.04 | 2.50E-08 | 0.69 | 0 | 0.6104/0.6042<br>0.6096/0.6042 | 0.58<br>(0.51, 0.65) |
| 19:10750738 | rs1560707 | upstream_gene | OA |  | T | G | 1.04 | 1.70E-13 | 1.02 | 1.91E-01 | 0.37 | 1.04 | 1.03 | 1.05 | 1.35E-13 | 0.45 | 0 | 0.3781/0.3683<br>0.382/0.3762 | 0.4<br>(0.33, 0.47) |
| 19:41833784 | rs75621460 | downstream_gene | OA | OA_kneehip | A | G | 1.16 | 4.40E-15 | 1.12 | 1.24E-02 | 0.03 | 1.16 | 1.12 | 1.2 | 1.62E-15 | 0.58 | 0 | 0.0205/0.0178<br>0.028/0.0252 | 0.03<br>(0.01, 0.07) |
| 19:55879672 | rs4252548 | missense | OA_hip |  | T | C | 1.37 | 5.30E-13 | 1.11 | 1.63E-01 | 0.02 | 1.32 | 1.22 | 1.43 | 1.96E-12 | 0.05 | 73 | 0.0276/0.0214<br>0.0258/0.0233 | 0.01<br>(0.002, 0.04) |
| 21:40048295 | rs2836618 | intergenic_region | OA_hip | OA_kneehip | A | G | 1.08 | 1.90E-07 | 1.18 | 2.17E-06 | 0.26 | 1.09 | 1.06 | 1.12 | 3.20E-11 | 0.02 | 82.6 | 0.2753/0.2609<br>0.2846/0.2524 | 0.19<br>(0.14, 0.26) |
| 22:43662241 | rs528981060 | upstream_gene | OA |  | A | G | 1.68 | 2.40E-08 |  |  | 0.001 | 1.68 | 1.4 | 2.02 | 2.37E-08 | 1 | 0 | 0.0012/0.0008<br>- | - |

| Previously reported loci |  |  |  |  |  |  |  |  |  |  |  |  |  |  |  |  |  |  |  |
| --- | --- | --- | --- | --- | --- | --- | --- | --- | --- | --- | --- | --- | --- | --- | --- | --- | --- | --- | --- |
| 1:219753509 | rs2820443 | intergenic_region | OA_kneehip | OA_hip;OA | C | T | 1.06 | 9.90E-10 | 1.06 | 1.20E-02 | 0.3 | 1.06 | 1.04 | 1.07 | 6.01E-11 | 0.82 | 0 | 0.3079/0.2968<br>0.312/0.2993 | 0.26<br>(0.2, 0.34) |
| 2:70717653 | rs3771501 | intron | OA | OA_hip;OA_kneehip | A | G | 1.05 | 6.00E-15 | 1.06 | 1.31E-02 | 0.47 | 1.05 | 1.03 | 1.06 | 4.24E-16 | 0.67 | 0 | 0.4838/0.4727<br>0.489/0.4753 | 0.48<br>(0.4, 0.55) |
| 3:52817778 | rs3774355 | upstream_gene | OA_hip | OA_kneehip | A | G | 1.09 | 3.00E-10 | 1.15 | 2.07E-05 | 0.36 | 1.09 | 1.07 | 1.12 | 8.20E-14 | 0.12 | 59 | 0.3776/0.358<br>0.3893/0.3574 | 0.39<br>(0.32, 0.47) |
| 6:45357699 | rs2396502 | intron | OA_hip | OA_kneehip | C | A | 1.09 | 1.80E-10 | 1.1 | 2.23E-03 | 0.6 | 1.09 | 1.06 | 1.11 | 2.12E-12 | 0.74 | 0 | 0.6218/0.6019<br>0.622/0.5997 | 0.64<br>(0.57, 0.71) |
| 6:76164589 | rs12209223 | upstream_gene | OA_hip |  | A | C | 1.16 | 2.50E-12 | 1.23 | 7.79E-06 | 0.1 | 1.17 | 1.13 | 1.21 | 3.88E-16 | 0.26 | 22 | 0.113/0.0998<br>0.1242/0.1036 | 0.09<br>(0.06, 0.15) |
| 9:4291928 | rs10974438 | intron | OA | OA_kneehip | A | C | 1.03 | 1.10E-07 | 1.06 | 6.39E-03 | 0.65 | 1.03 | 1.02 | 1.05 | 1.34E-08 | 0.36 | 0 | 0.6567/0.6494<br>0.658/0.6456 | 0.63<br>(0.55, 0.7) |
| 9:119484132 | rs34687269 | intron | OA_hip |  | A | T | 1.08 | 1.70E-10 | 1.1 | 1.67E-03 | 0.53 | 1.09 | 1.06 | 1.11 | 1.67E-12 | 0.7 | 0 | 0.5499/0.5299<br>0.5555/0.5323 | 0.52<br>(0.45, 0.6) |
| 12:28014970 | rs10492367 | intergenic_region | OA_hip | OA_kneehip | T | G | 1.16 | 2.90E-19 | 1.21 | 4.38E-07 | 0.19 | 1.16 | 1.13 | 1.2 | 1.25E-24 | 0.25 | 24.5 | 0.2093/0.187<br>0.2187/0.1877 | 0.18<br>(0.13, 0.25) |
| 15:58215727 | rs4775006 | intergenic_region | OA_knee |  | A | C | 1.05 | 1.30E-07 | 1.12 | 2.58E-04 | 0.41 | 1.06 | 1.04 | 1.08 | 8.40E-10 | 0.08 | 68.3 | 0.4227/0.41<br>0.437/0.4104 | 0.36<br>(0.29, 0.44) |
| 15:67370506 | rs12901372 | intron | OA_hip |  | C | G | 1.07 | 2.90E-08 | 1.13 | 5.22E-05 | 0.53 | 1.08 | 1.06 | 1.11 | 3.46E-11 | 0.13 | 56.2 | 0.5486/0.5319<br>0.5648/0.5347 | 0.47<br>(0.4, 0.55) |
| 16:53799977 | rs9930333 | intron | OA_kneehip |  | G | T | 1.04 | 4.30E-07 | 1.1 | 2.57E-04 | 0.42 | 1.05 | 1.03 | 1.06 | 1.52E-09 | 0.04 | 75.8 | 0.4316/0.4212<br>0.4471/0.4245 | 0.42<br>(0.35, 0.49) |
| 20:34025756 | rs143384 | 5_prime_UTR | OA_knee | OA_kneehip;OA | A | G | 1.1 | 3.60E-22 | 1.07 | 9.24E-03 | 0.6 | 1.1 | 1.08 | 1.12 | 4.77E-23 | 0.37 | 0 | 0.619/0.5959<br>0.6026/0.5865 | 0.64<br>(0.57, 0.71) |

Most Severe Consequence: Most severe consequence of index variant as predicted by SNPEff; Trait UKBiobank: Osteoarthritis trait most significantly associated with variant in the meta-analysis stage; Other Traits gwas sign metaanalysis: Other osteoarthritis traits with genome-wide significant association following meta-analysis; EA: Effect allele; NEA: Non-effect allele; OR: Odds ratio; PV: *P* value; WEAf: Weighted effect allele frequency between UK Biobank and arcOGEN; OR\_95L: Lower bound of the 95% credible interval of the odds ratio; OR\_95U: Upper bound of the 95% credible interval of the odds ratio; q\_pV: *P* value of Cochran's Q measure of heterogeneity; i2: $I^2$ statistic describing the percentage of variation across studies that is due to heterogeneity rather than chance; OA: osteoarthritis; OA\_hip: Hip osteoarthritis; OA\_knee: Knee osteoarthritis; OA\_kneehip: Knee and/or hip osteoarthritis; \* UK Biobank allele frequencies are displayed at the top with arcOGEN below. \*\*1000 Genomes Project GBR lower and upper 95% confidence intervals for the EAF are displayed in brackets.

To identify putative effector genes at the 64 genome-wide significant regions, we integrated results from several strands of investigation, including transcriptomic/proteomic characterisation of primary tissue from osteoarthritis patients undergoing joint replacement surgery, coupled with statistical fine-mapping, annotation of predicted consequences of variants in the credible sets, eQTL colocalization, and relevant rare human disease and animal model evidence (Online Methods; Supplementary Table 4 and 5). We observe evidence of colocalization in at least one tissue for 49 out of the 64 loci, 44 of which are at newly-associated osteoarthritis signals (Supplementary Table 6). Using MetaXcan, we identify 11 genes with additional evidence of colocalization at loci not reaching genome-wide significance in SNV analyses (Supplementary Figure 7; Supplementary Tables 7-8).

Pathway analyses (Online Methods, Supplementary Note) identify 64 biological processes associated with osteoarthritis, of which 46 are bone-, cartilage‐ and chondrocyte‐ morphology related (Supplementary Table 9). The collagen formation and extracellular matrix organisation biological pathways are consistently identified by different pathway analysis methods. Genome-wide linkage disequilibrium (LD) score regression analysis^5,6^ unveils significant correlation between osteoarthritis and traits within the obesity, cognition, smoking, bone mineral density and reproductive trait categories (Figure 1; Supplementary Table 10 and 11). Mendelian randomization analyses (Online Methods) support a role for higher body mass index (BMI) and adiposity in osteoarthritis risk, and identify a potential protective effect of LDL cholesterol, and of higher level of education against osteoarthritis (Supplementary Tables 12-14, Supplementary Note). Two of the BMI loci (*SLC39A8* and *FTO*) show genome-wide significant associations with osteoarthritis, with *SLC39A8* showing much larger effects on osteoarthritis than expected given the BMI-raising effects (Supplementary Figure 8). Apparent causal associations of knee pain with osteoarthritis (Supplementary Table 12 and 15) are potentially attributable to reverse causality (Supplementary Note). We estimate the proportion of the total narrow sense heritability explained by osteoarthritis loci to be 14.7 % for knee osteoarthritis, 51.9 % for hip osteoarthritis, 24.2% for osteoarthritis of the hip and/or knee, and 22.5% of osteoarthritis at any site (Supplementary Table 16). We do not find evidence for a role of low-frequency or rare variation of large effect in osteoarthritis susceptibility, and have limited power to detect smaller effects at lower-frequency variants (Figure 2). In the future, meta-analyses of osteoarthritis studies in global populations will help further deconvolute the genetic underpinning of this disabling disease.

**Figure 1:**
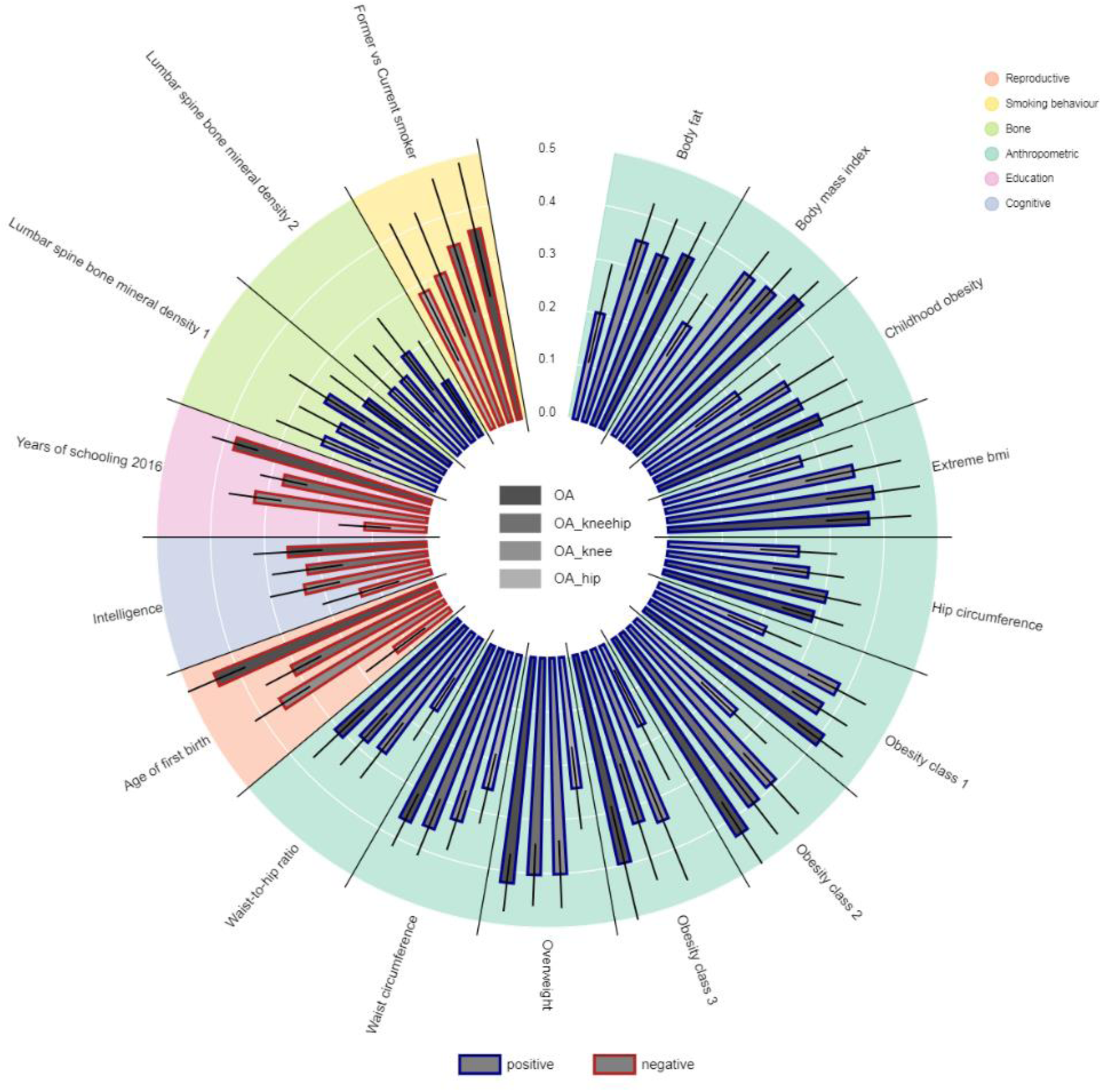
Genetic correlations between osteoarthritis and other traits and diseases. Genetic correlations (rg) between osteoarthritis and other publicly available GWAS results, based on LD score regression as implemented in LDHub. The diagram shows traits with significant correlation (*P*<0.05) and 95% confidence intervals across all osteoarthritis definitions. The red outline of the bars denotes negative correlation and the blue outline denotes positive correlation. The upper right legend shows the categories of the traits. OA: osteoarthritis; OA_hip: Hip osteoarthritis; OA_knee: Knee osteoarthritis; OA_kneehip: Knee and/or hip osteoarthritis. Lumbar spine bone mineral density 1 and 2 relate to two different published studies.

**Figure 2:**
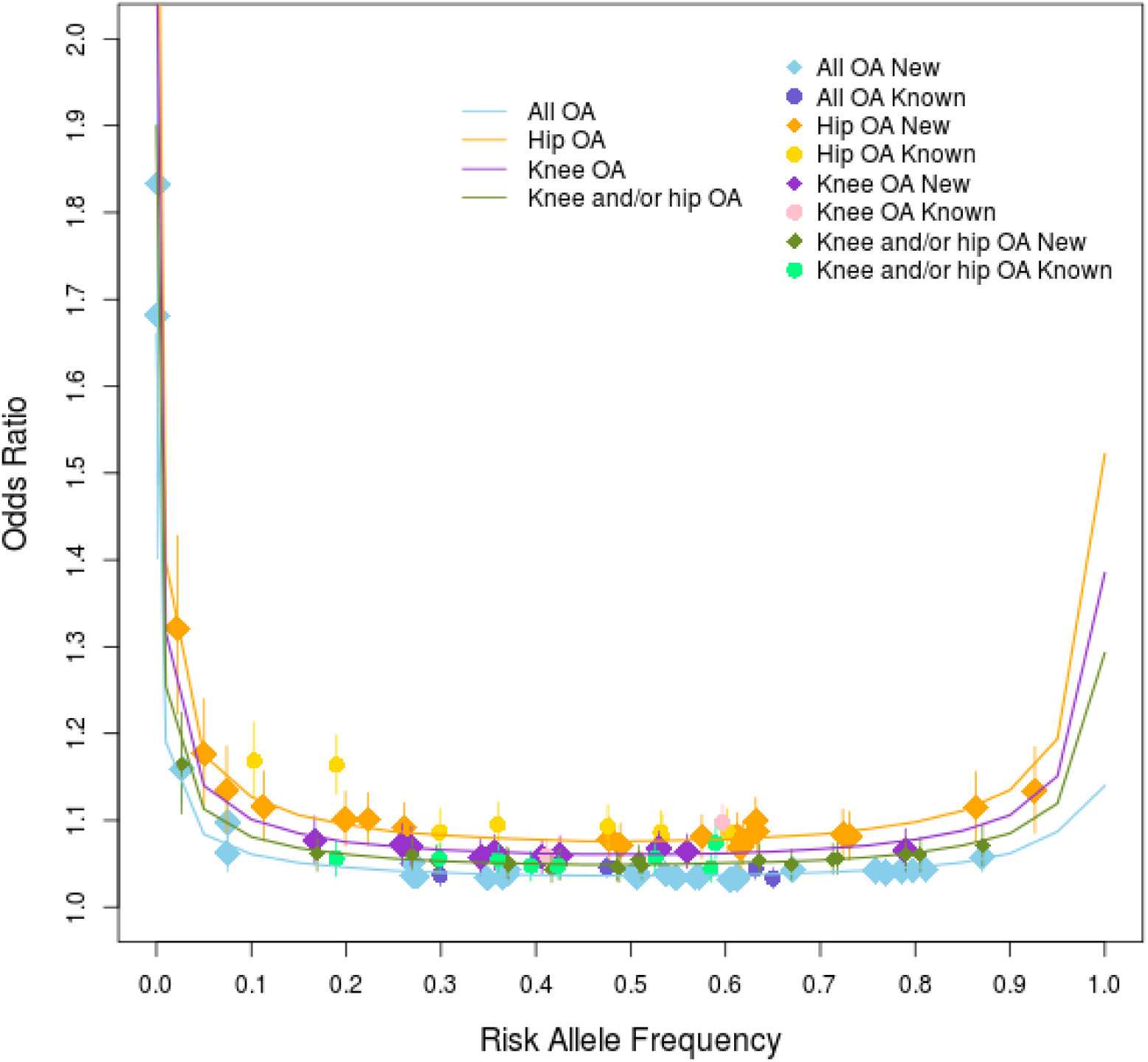
Allelic architecture of index variants. Meta-analysis based odds ratio with its 95% confidence interval of 99 variants (previously-reported denoted as circles and newly-reported denoted as diamonds) with UK Biobank and arcOGEN meta-analysis *P*<3.0×10^-8^ (two-sided) as a function of their weighted allele frequency. The curves indicate 80% power at the genome-wide significance threshold of *P*≤3.0×10^-8^, for the four sample sizes of the meta-analyses. We have 80% power to detect an association at genome-wide significance for a variant with 1% MAF and allelic odds ratio of 1.19, 1.40, 1.32 and 1.25 for all osteoarthritis, hip osteoarthritis, knee osteoarthritis and knee and/or hip osteoarthritis, respectively. For 0.1% MAF the corresponding odds ratios are 1.66, 2.43, 2.12 and 1.90.

We used a combination of conditional analyses^7^ followed by asymptotic Bayes’ factor fine-mapping^8^ (Online Methods) of conditionally distinct association signals to identify causal variants. In six of the novel loci, a single variant could be postulated as causal with more than 95% posterior probability: missense variants in *SLC39A8*, *IL11* and *ANAPC4* (rs13107325, rs4252548 and rs34811474, respectively), rs75621460 near *TGFB1*, rs547116051 near *MAPT* and rs528981060 near *SCUBE1* (Supplementary Table 17, Supplementary Note).

We observe strong enrichment for genes known to cause monogenic bone development diseases and forms of early-onset osteoarthritis, in the vicinity of osteoarthritis signals (odds ratio [OR] 8.87, *P*=1.8×10^-4^, and OR 8.83, *P*=8×10^-3^, respectively) (Supplementary Table 18 and 19). This finding highlights bone development as an important physiological process in osteoarthritis aetiology. Several genes identified as likely causal in our study are also linked to osteoarthritis aetiology in animal models. In eight out of the ten cases where we can unequivocally define directionality of association, we observe concordance between our results and those from animal models (i.e. that reduced expression or loss-of-function mutations increase osteoarthritis risk both in humans and in animal models) (Supplementary Table 20). Some of these genes code for structural bone/cartilage proteins (*COL11A1*, *COL11A2*) or play a critical role in bone/cartilage development (*FGFR3*, *GDF5*). These consistent observations in human and animal models provide compelling evidence for a causal role of these genes in osteoarthritis and point to an agonist strategy as the desired mechanism of action for new osteoarthritis drugs targeting these eight genes.

Ten genes have a therapeutic approved or in clinical trials (Table 2), with mechanisms of action that are not inconsistent with potential for efficacy in osteoarthritis, based on eQTL, functional genomics, rare disease and animal model data. Four of these genes, *TGFB1*, *GDF5*, *FGF18* and *CTSK*, currently have therapeutics in clinical development for osteoarthritis/cartilage regeneration indications. Of these, only *GDF5* has been previously published as genetically associated with osteoarthritis susceptibility^9^. Two of the genes, *IL11* and *DPEP1*, have approved therapeutics for unrelated indications, opening the possibility for repositioning.

**Table 2:**
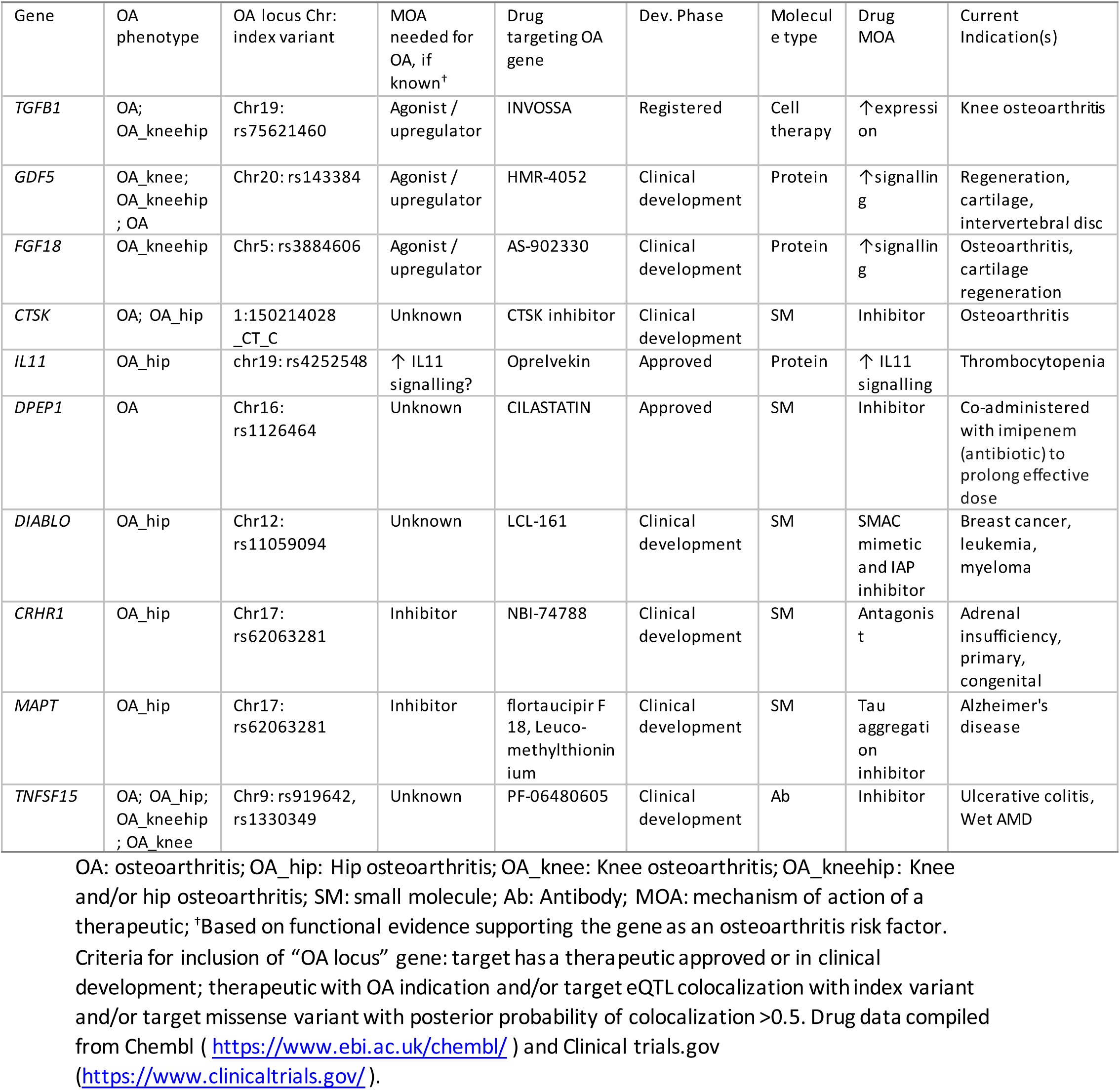
**Translational context for selected osteoarthritis-associated genes.**

rs4252548 (hip osteoarthritis, posterior probability of causality [PPC] 0.99), is a predicted deleterious missense variant (Arg112His) in *IL11* (interleukin 11), associated with increased risk of hip osteoarthritis. Using RNA sequencing (Online Methods), we find that *IL11* shows increased expression in degraded compared to intact cartilage (log2 fold change [logFC]=0.787, false discovery rate [FDR]=4.82×10^-3^). This cytokine is a potent stimulator of bone formation^10^, is required for normal bone turnover^11^ and has been previously found to be up-regulated in osteoarthritis knee tissue and to be associated with disease progression^12^. The rs4252548 osteoarthritis risk allele is also associated with decreased adult height^13^. A recombinant human IL11 molecule (NEUMEGA) with three-fold-enhanced affinity for IL11RA, compared to IL11^14^, is approved for the treatment of chemotherapy-induced thrombocytopenia (Table 2). The likely effects of increased IL11 signalling in osteoarthritis joints are currently not well understood, and it is worth evaluating this therapeutic for potential efficacy in disease models.

The rs1126464 (osteoarthritis, PPC 0.89) locus index signal is a missense variant (Glu351Gln) in *DPEP1*, predicted to be tolerated. *DPEP1* hydrolyses a wide range of dipeptides, and is implicated in the renal metabolism of glutathione and its conjugates. A DPEP1 inhibitor, cilastatin, is approved and used in combination with the antibiotic imipenem, in order to protect it from dehydropeptidase and prolong its antibacterial effect^15^. We suggest investigating the effects of cilastatin in osteoarthritis models to determine whether this has potential as a therapeutic, or whether an agonist may be efficacious.

rs75621460 (hip and/or knee osteoarthritis, single variant in the 95% credible set) is an intergenic variant residing downstream of *CCDC97* and *TGFB1* (Table 1), and is colocalised with a *TGFB1* eQTL in sun-exposed skin (GTEx) (Supplementary Table 6). Mutations in *TGFB1* cause Camurati–Engelmann disease characterised by diaphyseal dysplasia with thickening and fluctuating bone volume giving rise to bone pain, muscle weakness, gait issues and tiredness^16,17^. *TGFB1* plays a critical role in skeletal development and adult bone homeostasis^18^, including bone remodelling^19^, osteoclast/osteoblast differentiation^20,21^ and chondrogenesis^22^. INVOSSA^TM^, a TGFB1 cell and gene therapy in chondrocytes, was associated with significant improvements in function and pain in patients with knee osteoarthritis^23^.

The importance of TGFB1 signalling for osteoarthritis is supported by significant enrichment for “TGF Beta Signalling Pathway” genes (Supplementary table 9), including: *LTBP1*, *LTBP3*, *SMAD3* and *RUNX2*. *LTBP1*, at the novel rs4671010 locus, directly interacts with *TGFB1*, is involved in the assembly, secretion and targeting of *TGFB1* to sites at which it is stored and/or activated, and may contribute to controlling the activity of *TGFB1*^24^. *LTBP3* (novel locus: rs10896015) directly interacts with and activates *TGFB1* in the early proliferative phase of osteogenic differentiation^25^. *SMAD3* (known locus: rs12901372) plays a critical role in chondrogenic differentiation, and regulates *TGFB1* expression^26^; and the *TGFB1/SMAD3* pathway regulates the expression of miR-140 in osteoarthritis^27^. The directionality of the colocalized eQTL and animal model data suggest that agonism/up-regulation of LTBP1, LTBP3 and SMAD3 may be therapeutic for osteoarthritis (Supplementary Table 20). *RUNX2* (known locus: rs2064630) is a transcription factor essential for the osteoblast differentiation and chondrocyte maturation^28^, and is down-regulated by *TGFB1*^29^. Given the genetic and biological support for the importance of *TGFB1* in osteoarthritis aetiology and treatment, there may be scope for the development of simpler osteoarthritis therapeutics which target this mechanism, such as a small molecule or antibody.

Although not a current drug target, the novel *SLC39A8* association is noteworthy. rs13107325 (osteoarthritis, PPC 0.99) is a missense variant located in *SLC39A8* and demonstrates significantly increased expression in degraded compared to intact articular cartilage (logFC=0.522, FDR=5.80×10^-5^) (Table1; Supplementary Table 2), consistent with previously-reported increased levels of *SLC39A8* in osteoarthritis compared to healthy chondrocytes^30,31^. rs13107325 is also associated with obesity^32^, hypertension^33^, Crohn’s disease and altered microbiome composition^34^. *SLC39A8* functions in the cellular import of zinc at the onset of inflammation. Suppression of *SLC39A8* has been shown to reduce cartilage degradation in osteoarthritis animal models^30^. The zinc‐ *SLC39A8*-MTF1 axis has been proposed to be an essential catabolic regulator of osteoarthritis pathogenesis^31^.

In this study, we have more than doubled the number of osteoarthritis risk loci, supported by integrated eQTL colocalization, fine-mapping, Mendelian bone disease, animal model and differential osteoarthritis joint expression data, to reveal putative effector genes. In addition to identifying chondrocyte and osteoblast biological mechanisms implicated in osteoarthritis susceptibility, we have revealed biological mechanisms that represent attractive targets for osteoarthritis drug discovery, and highlight approved therapeutics which represent viable considerations for repositioning as osteoarthritis therapies. We anticipate that this advance in basic understanding of osteoarthritis risk factors and mechanisms will stimulate the evaluation of novel drug targets for osteoarthritis.

## ONLINE METHODS

### Studies

*UK Biobank*: UK Biobank is a cohort of 500,000 participants aged 40-69 years recruited between 2006 and 2010 in 22 assessment centres throughout the UK^35^. The assessment visit included electronic signed consent; a self-completed touch-screen questionnaire; brief computer-assisted interview; physical and functional measures; and collection of biological samples and genetic data. This work was based on the third UK Biobank release, which includes the full set of the 500,000 genotypes imputed on the Haplotype Reference Consortium^36^ and the 1000 Genomes Consortium^37^. Briefly, 50,000 samples were genotyped using the UKBiLEVE array and the remaining samples were genotyped using the UK Biobank Axiom array (Affymetrix). After sample and SNP quality control (QC) of the directly-typed genotypes, followed by phasing and imputation, carried out centrally^38^, there are approximately 97million variants in 487,411 individuals. Following additional QC checks, we excluded samples with call rate ≤95%. We checked samples for gender discrepancies, excess heterozygosity, over third degree relatedness, ethnicity and we removed possibly contaminated and withdrawn samples. Variants with minor allele frequency (MAF) of ≤0.1% or effective minor allele count (imputation quality score times minor allele count) ≤90 were discarded. In summary, we interrogated approximately 17 million variants from 450,805 related individuals of European descent (Supplementary Figure 1).

To define osteoarthritis cases, we used the self-reported status established during interview with a nurse (Field 20002; Initial assessment visit) and the Hospital Episode Statistics ICD10 primary and secondary codes. We conducted four osteoarthritis discovery GWAS: self-reported or hospital-diagnosed osteoarthritis at any site based on ICD10 hospital record codes M15-M19 (n=70,532); hospital-diagnosed hip osteoarthritis based on ICD10 hospital record M16 (n=12,850); hospital-diagnosed knee osteoarthritis based on ICD10 hospital record M17 codes (n=21,921); and hospital-diagnosed hip and/or knee osteoarthritis (M16 or M17; n=32,907). To minimise misclassification in the control datasets to the extent possible, in all analyses, from the controls, we excluded individuals with primary or secondary ICD10 codes M05 through M14, as well as arthrosis codes M15-M19. This excluded individuals with inflammatory polyarthropathies and resulted in a control sample size of 369,983 controls (Supplementary Figure 1).

Principle components (PCs) were computed using fastPCA^39^ and high quality directly typed markers from the unrelated set of Europeans. PCs were then projected to the related European sample using SNP weights. We tested for association using the non-infinitesimal mixed model association test implemented in BOLT-LMM v2.3^40^ with adjustment for the first 10 PCs, sex, age at recruitment and genotyping chip. As BOLT-LMM implements a linear regression, effect size estimates and their standard errors for case-control outcomes have been calculated by dividing the effect size estimates and standard errors output by the software by *p* × (1 - *p*), where *p* is the proportion of cases in the trait definition^41^.

*Arthritis Research UK Osteoarthritis Genetics (arcOGEN) – cases*: arcOGEN is a collection of unrelated, UK-based individuals of European ancestry with knee and/or hip osteoarthritis from the arcOGEN Consortium^9,42^. Cases were ascertained based on clinical evidence of disease to a level requiring joint replacement or radiographic evidence of disease (Kellgren–Lawrence grade ≥2). The arcOGEN study was ethically approved, and all subjects used in this study provided written, informed consent.

*United Kingdom Household Longitudinal Study (UKHLS) – controls*: The UKHLS, also known as Understanding Society, is a longitudinal panel survey of 40,000 UK households (England, Scotland, Wales and Northern Ireland) representative of the UK population. Participants are surveyed annually since 2009 and contribute information relating to their socioeconomic circumstances, attitudes, and behaviours via a computer assisted interview. The study includes phenotypical data for a representative sample of participants for a wide range of social and economic indicators as well as a biological sample collection encompassing biometric, physiological, biochemical, and haematological measurements and self-reported medical history and medication use. The UKHLS has been approved by the University of Essex Ethics Committee and informed consent was obtained from every participant.

*Genotyping, imputation and association testing*: 7,410 arcOGEN cases were genotyped on the Illumina Human 610-Quad array, 670 arcOGEN cases were genotyped on the Illumina OmniExpress array, and 9,296 UKHLS controls were genotyped on the Illumina CoreExome array. Genotype QC has been previously described^9,43,44^. Prior to imputation all variants were mapped to GRCh37 and cases and controls were merged into a single dataset containing overlapping variants. We further excluded variants with MAF≤1, indels, and evidence for differential missingness between cases and controls (Fisher’s exact test *P*<0.0001). We performed a case-control analysis and visually inspected the cluster plots for any variant with *P*≤5×10^-8^; variants with poor clusters were excluded. Final QC checks prior to imputation were performed using a HRC pre-imputation checking tool (URLs). Imputation was performed on 17,376 individuals and 126,188 variants using the Michigan HRC^36,45^ imputation server with Eagle2^46^ for the prephasing. We excluded related individuals by performing pair-wise identity-by-descent (IBD) (using PI_HAT threshold > 0.2) in PLINK^47^ using directly typed variants with MAF>1% and pruned for LD using r^2^<0.2. In addition, we excluded any individuals that were related to individuals in the UK Biobank dataset using the imputed variants and the same IBD method as described above. We also excluded any variant that had an imputation information score (minimac R-square)<0.3, with a Hardy Weinberg *P*≤1×10^-6^ and MAF ≤ 0.1%The final dataset contained ∼11.6 million variants in 6,520 cases and 8,186 controls. Association analysis was performed using SNPTEST^48^ with the first ten principal components as covariates. The arcOGEN dataset comprised three osteoarthritis phenotypes: hip osteoarthritis (2,854 cases and 8,186 controls), knee osteoarthritis (3,034 cases and 8,186 controls), and hip and/or knee osteoarthritis (6,520 cases and 8,186 controls).

### Meta-analysis

We meta-analysed the UK Biobank and arcOGEN datasets using fixed effects inverse-variance weighted meta-analysis in METAL^49^. We performed meta-analyses across osteoarthritis definitions using summary statistics from the UK Biobank and arcOGEN cohorts, and defined genome-wide significance based on the meta-analysis combined *P* value as outlined below.

### Significance threshold

The osteoarthritis traits analysed in this study are highly correlated. To calculate *M*_*eff* the effective number of independent traits, we estimated the genetic correlation matrix between the 4 osteoarthritis traits (Supplementary Table 1) using LDscore^50^ with genome-wide summary statistics of common-frequency variants in the UK Biobank dataset. We then calculated *M*_*eff* from the eigenvalues *λ*_*i* of the correlation matrix^51^:

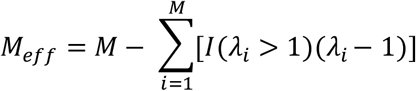

For the *M* = *4* osteoarthritis phenotypes in this study, *M*_*eff* = *1*.*6046*. We therefore use *P*≤3×10^-8^ as the threshold corrected for the effective number of traits to report genome wide significance.

### Statistical independence

To define independent signals within a GWAS, we performed physical clumping using a simple iterative procedure. We rank all variants that reach a *P* value threshold according to their *P* value. The variant with the smallest *P* value is considered the index variant of that signal and any variants within 1MB region either side of that index variant that reach the pre-defined *P* value threshold are clumped with that variant. We repeat the procedure until no more variants that reach the predefined *P* value threshold exist that have not been assigned to a physical clump. To test that the index variants defined by this procedure are statistically independent, we performed an approximate stepwise model selection procedure, as implemented by COJO in GCTA^7^. An independent signal in a region is declared if its *P* value of association in the stepwise regression is less than 3×10^-8^. LD calculations were based on the full UK Biobank imputed set.

To define independent signals across the four osteoarthritis GWAS, we performed reciprocal approximate conditional analyses, as implemented by COJO in GCTA^7^, of each index variant of one GWAS conditioned on each index variant of the other GWAS. A signal between two GWAS is considered to be the same if the *P* value of an index variant of one GWAS conditioned on an index variant of the other GWAS is ≤ 10-5 or a *P* value difference between conditional and unconditional analysis of less than 2 orders of magnitude.

To investigate statistical independence between index variants from each GWAS and previously reported variants, we performed approximate conditional analysis, as implemented by COJO in GCTA^7^, of each index variant conditional on all previously reported variants within 1Mb region, each one at a time. The index variant was considered independent from a previously reported variant if it had a conditional *P* ≤ 10^-5^ or a *P* value difference between conditional and unconditional analysis of less than 2 orders of magnitude. Variants were classified as known (denoting either a previously reported variant, or a variant for which the association signal disappears after conditioning on the lead variant of a previously reported locus) or newly identified (denoting a variant which is conditionally independent of previously reported loci).

### Fine-mapping

We constructed regions for fine-mapping by taking a window of 1Mb either side of each index variant. Within each region, we performed an approximate stepwise model selection procedure, as implemented by COJO in GCTA^7^, using the meta-analysis summary statistics and LD calculations based on the UK Biobank cohort to determine the number of independent signals. We consider conditionally distinct signals those where the stepwise regression association reaches genome-wide significance (*P*<3.0×10^-8^). We then perform single-SNP approximate association analyses conditional on the set of SNPs identified by the model selection procedure, again using COJO, and we calculate Wakefield’s asymptotic Bayes’ factors^8^ (ABF). In particular, when there is a single causal variant in the region, ABF is based on the marginal summary statistics of the meta-analysis. When there are multiple causal variants in the region, for each signal we calculate a set of ABF using the conditional summary statistics of the meta-analysis conditioned on all other signals. For each signal, we then calculate posterior probabilities of each variant being causal and a 95% credible set, which contains the minimum set of variants that jointly have at least 95% probability of including the causal variant. As this number can be large, we focus on the variants in the 95% credible set that have posterior probability of causality (PPC) over 3% and also on any variants in the 95% credible set with moderate or high consequence (irrespective of their PPC).

### Genetic correlation analysis

To better understand the degree to which genetic architecture is shared across osteoarthritis and other complex traits, LD score regression^5^ was performed as implemented in the LDHub pipeline^6^ (URLs). We calculated the genome-wide genetic correlation between each of the osteoarthritis definitions and all available 832 human traits and diseases (accessed 15-18 June 2018). Of these, 597 traits were available within the UK Biobank resource. In each analysis, all variants in the major histocompatibility complex (MHC) region on chromosome 6 (26–34 MB) were removed and only variants with rsIDs were included in the analyses, yielding 1203892 - 1204029 variants overlapping with LDHub. We used the Benjamini-Hochberg false discovery rate and the effective number of independent traits tested for multiple testing correction. The level of significance was set at FDR-corrected *P*<0.05.

### Mendelian randomization

We performed Mendelian randomization analyses using the MR-Base platform^52^. We tested the bidirectional causal associations of each of the four osteoarthritis datasets with 991 exposures/ outcomes in MR-Base. Statistical significance was considered at *P*<6.3×10^-6^. To follow up on pain associations, we performed analyses of knee and hip pain as an outcome after excluding all individuals self-reporting or hospital-diagnosed with osteoarthritis in UK Biobank. All instruments were aggressively clumped prior to analysis (LD r^2^<0.001) and inverse variance-weighted (IVW), Median-weighted, and MR-Egger analyses were performed for multi-variant instruments, and Wald ratio estimators were used to assess causality for single variant instruments.

### Transcriptome-wide association

We used a gene-based approach, MetaXcan^53^, to test for associations between the osteoarthritis traits and predicted expression levels in 48 human tissues from GTEx V7^54^. MetaXcan leverages a set of reference individuals for whom both gene expression and genetic variation have been measured to impute the cis-genetic component of expression into a much larger set using the elastic net model. It then correlates the imputed gene expression to the trait of interest and performs a transcriptome-wide association study to identify significant expression-trait associations. We used a conservative Bonferroni correction to account for the gene-tissue pairs (20,000 genes across 48 tissues), leading to a significance threshold of 5.20×10^-8^. To reduce the effect of LD confounding on the MetaXcan results, when different causal SNPs are affecting expression levels and the phenotypic trait in a GWAS, we estimated the probability of colocalization of each GWAS and expression quantitative trait locus (eQTL) signal in each significant MetaXcan result using Coloc^55^ (Supplementary Note; Supplementary Tables 7-8; Supplementary Figure 7).

### Colocalization analysis

To assess whether the genome-wide significant osteoarthritis signals colocalise with eQTL signals, and therefore potentially share a causal molecular mechanism, we employed the Coloc method^55^, which uses asymptotic Bayes factors with summary statistics and regional LD structure to estimate five posterior probabilities: no association with either GWAS or eQTL (PP0), association with GWAS only (PP1), association with eQTL only (PP2), association with GWAS and eQTL but two independent SNPs (PP3), and association with GWAS and eQTL having one shared SNP (PP4). A large posterior probability for PP4 indicates support for a single variant affecting both GWAS and eQTL studies. For each of the GWAS signals, we defined a 100kb region either side of the index variant, and tested for colocalization within the entire cis-region of any overlapping eQTLs (transcription start and end position of an eQTL gene plus and/or minus 1Mb, as defined by GTEx) in 48 human tissues from GTEx V7^54^. A PP4 over or equal to 80% was considered as evidence for colocalization (Supplementary Note; Supplementary Table 6).

Most colocalization methods, such as Coloc, rely on the availability of genome-wide eQTL results, which are not always readily available. For eQTL datasets with no publically available full summary statistics, we used an alternative approach that estimates the probability of colocalization using published top eQTL signals. First, we estimated the credible sets for the eQTLs using the Probabilistic Identification of Causal SNPs (PICS) method^56^ for each index SNP for each gene from 27 eQTL studies (Supplementary Table 5). PICS is a fine-mapping algorithm that assumes one causal signal tagged by a single index SNP per locus. For neutral SNPs (SNPs whose association signals are due to LD with the causal SNP), the strength of association scales linearly with the r^2^ relationship/distance to the index SNP. Under this assumption, PICS can estimate the posterior probability of a given SNP being causal using LD information from the 1000 Genomes database. Second, we generated PICs credible sets for osteoarthritis GWAS index SNPs. We then performed a colocalization analysis of the osteoarthritis GWAS and eQTL PICs credible sets using an adapted Coloc method^50^. Given that PICs calculates the posterior probabilities for each SNP in the credible set, we bypassed the need for calculating the Bayes Factors using Wakefield’s approximate Bayes Factor method which is reliant on full summary statistics. Colocalizations with a posterior probability greater than 0.8 were considered positive. This method was benchmarked on other GWAS datasets, and we found the false positive rate to be no higher than the standard Coloc package.

We observe evidence of colocalization in at least one tissue for 50 out of our 64 loci using any of the 3 methods (MetaXcan, Coloc, Piccolo), 41 of which are at newly associated osteoarthritis signals (Supplementary Table 6). MetaXcan alone identified 119 genes, Coloc 113 and Piccolo 58, while the overlap of all 3 methods implicate 20 genes (*TGFA, ILF3, CSK, CYP1A1, ULK3, CHMP1A, TSKU, SUPT3H, GNL3, NT5DC2, LMX1B, SMAD3, MLXIP, COLGALT2, FAM89B, UQCC1, NFAT5, ALDH1A2, FAM53A, FGFR3*; Supplementary Figure 9).

### Heritability estimation

To investigate the narrow sense heritability for the four osteoarthritis disease definitions, we ran LDscore^50^, which uses summary statistics at common-frequency variants genome-wide (independent of *P* value thresholds) and LD estimates between variants while accounting for sample overlap. To calculate the population prevalence in the UK (65 million people), we consulted Arthritis Research UK figures: 8.75 million people have symptomatic osteoarthritis, while 2.46 and 4.11 million people have osteoarthritis of the hip and the knee, respectively. We assumed that 2.46+4.11 million people have osteoarthritis of the hip and/or the knee. We estimated the phenotypic variance explained by the 99previously and newly reported variants that reached genome-wide significance in the meta-analysis between UK Biobank and arcOGEN, as a function of allele frequency (Figure 2; Supplementary Table 18). The phenotypic variance explained by a variant is) *ln*(*OR*)^2^ × 2 × *EAF* × (1 – *EAF*), where ln(*OR*) is the natural logarithm of the OR of the variant in the meta-analysis and *EAF* is its weighted effect allele frequency across UK Biobank and arcOGEN. Variants associated with hip osteoarthritis tend to have larger effect size estimates and hence explain more of the phenotypic variability (Figure 2; Supplementary Table 18). The hip osteoarthritis dataset is the smallest in both the UK Biobank and arcOGEN cohorts (18% and 59% fewer cases compared to knee osteoarthritis and osteoarthritis at any joint in UK Biobank, respectively).

### Pathway analysis

We performed gene-set analyses for each of the osteoarthritis phenotypes separately, using MAGMA v1.06^57^. We mapped variants to 19,427 protein-coding genes (NCBI 37.3), including a 10kb window on either side of the gene. We then computed gene *P* values based on individual variant association *P* values. We used the ‘snp-wise=mean’ model, which calculates the mean of the χ^2-^statistic amongst the single variant *P* values in each gene, and applied default MAGMA QC steps. Genotype data of 10,000 individuals (subset of self-reported plus hospital-diagnosed osteoarthritis at any site analysis), were used to calculate LD (as measured by r^2^). We carried out a one-sided competitive gene-set analysis for each phenotype, implemented as a linear regression model on a gene data matrix created internally from the gene-based results. Briefly, this converts the gene-based *P* values to Z-scores, and tests if the mean association with the phenotype of genes in the gene set is greater than that of all other genes. We used Kyoto Encyclopedia of Genes and Reactome (accessed through MSigDB113 (version 5.2) on 23 January 2017). We also downloaded Gene Ontology (GO) biological process and molecular function gene annotations from Ensembl (version 87). We used annotations with the following evidence codes: a) Inferred from Mutant Phenotype (IMP); b) Inferred from Physical Interaction (IPI); c) Inferred from Direct Assay (IDA); d) Inferred from Expression Pattern (IEP); and e) Traceable Author Statement (TAS). KEGG/Reactome and GO annotations were analysed separately and only pathways that contained between 20 and 200 genes were included (594 for KEGG/Reactome, 619 for GO). We used MAGMA’s built-in permutation method (k=10,000 permutations) to produce corrected competitive *P* values with a family-wise error rate (FWER) of 5%. We then further adjusted these corrected competitive *P* values for the effective number of independent traits tested (1.6046).

We also performed gene set enrichment analysis by using DEPICT (URLs) and PASCAL (URLs). DEPICT version 1 rel194 was downloaded from GitHub (URLs) on 14/06/2018. We run DEPICT separately in each of the four osteoarthritis definitions for the variants with a meta-analysis *P*<1×10^-5^. Briefly, DEPICT first clumped the variants with *P*<1×10^-5^ using 500 kb flanking regions as physical distance threshold and an r^2^>0.1 with PLINK^47^ to obtain lists of independent SNPs, resulting in 864 clumps. Variants within the major histocompatibility complex region on chromosome 6 were excluded. DEPICT analyses were conducted using the default settings: 50 repetitions to compute FDR and 500 permutations based on 500 null GWAS to compute *P* values adjusted for gene length. All 14,461 available reconstituted gene sets were used representing a wide spectrum of biological and mouse phenotypic annotations. We also used the method implemented in PASCAL to perform gene set enrichment analysis which accounts for LD structure in the genome and particularly of highly correlated chromosomal regions containing multiple genes that can negatively impact the results of the analysis. In this approach, variants were first mapped to genes, including a 10kb window on either side of the gene. We then computed gene scores by aggregating the single-marker association values with the LD structure. Finally, the scores of genes that belong to the same pathways (i.e. gene sets) were used to compute pathway scores and determine the statistical significance of the association between the pathway and each of the osteoarthritis phenotypes. Here we used exactly the same pathways of the MAGMA analysis. The gene and the pathway scores were performed by using the sum gene score and the chi-squared approach respectively, as implemented in PASCAL. All pathway *P* values obtained by either software were adjusted for multiple testing correction by using FDR and the effective number of independent traits. The level of significance was set at FDR-corrected *P*<0.05.

### Monogenic enrichment analysis

We compiled a systematic list of genes causing bone phenotypes in humans by scanning the STOPGAP database^53^, which uses OMIM (URLs) and Orphanet (URLs) to define genes underlying monogenic/Mendelian diseases. We selected all genes causing monogenic diseases and annotated with MeSH terms (Medical Subject Headings) related to bone, cartilage or joint disease, including: “bone disease, developmental”, “osteochondrodysplasias”, “osteogenesis imperfecta”, “osteoporosis”, “osteopetrosis”, “arthritis, juvenile” and “arthrogryposis”. Other bone-, cartilage or joint related mesh terms linked to less than 10 genes in the STOPGAP database were excluded from the analysis. Additionally, we selected a list of well-validated genes underlying syndromic or nonsyndromic forms of early onset osteoarthritis (EO-OA) from a review by Aury-Landas et al.^54^. For enrichment analysis, genes residing within 500kb of each index variant identified in our GWAS were considered as osteoarthritis loci, and the rest of the genes in the genome associated to any mesh term in STOPGAP were considered non-osteoarthritis loci. We built a 2×2 table by counting the number of genes annotated to each of the above-mentioned MeSH terms among osteoarthritis and non-osteoarthritis loci. We assessed evidence for enrichment using a Fisher’s exact test.

### Transcriptomic and proteomic analyses

*Patients and samples*: We collected cartilage samples from 38 patients undergoing total joint replacement surgery: 12 knee osteoarthritis patients (cohort 1; 2 women, 10 men, age 50-88 years); knee osteoarthritis patients (cohort 2; 12 women 5 men, age 54-82 years); 9 hip osteoarthritis patients (cohort 3; 6 women, 3 men, age 44-84 years). We collected matched intact and degraded cartilage samples from each patient. Cartilage was separated from bone and chondrocytes were extracted from each sample. From each isolated chondrocyte sample, we extracted DNA, RNA and protein. All patients provided full written informed consent prior to participation. The human biological samples were sourced ethically and their research use was in accord with the terms of the informed consents under an IRB/EC approved protocol. All sample collection, DNA, RNA and protein analysis steps are described in detail in Steinberg et al^58^.

*Proteomics*: Proteomics analysis was performed on intact and degraded cartilage samples from 24 individuals (15 from cohort 2, 9 from cohort 3). LC-MS analysis was performed on the Dionex Ultimate 3000 UHPLC system coupled with the Orbitrap Fusion Tribrid Mass Spectrometer. To account for protein loading, abundance values were normalised by the sum of all protein abundances in a given sample, then log2-transformed and quantile normalised. We restricted the analysis to 3917 proteins that were quantified in all samples. We tested proteins for differential abundance using limma^59^ in R, based on a within-individual paired sample design. Significance was defined at 1% Benjamini-Hochberg FDR to correct for multiple testing. Of the 3732 proteins with unique mapping of gene name and Ensembl ID, we took forward 245 and 489 proteins with significantly different abundance between intact and degraded cartilage at 1% and 5% FDR, respectively (Supplementary table 2).

*RNA sequencing*: We performed a gene expression analysis on samples from all 38 patients. Multiplexed libraries were sequenced on the Illumina HiSeq 2000 (75bp paired-end read length). This yielded bam files for cohort 1 and cram files for cohorts 2 and 3. The cram files were converted to bam files using samtools 1.3.1^60^ and then to fastq files using biobambam 0.0.191^61^, after exclusion of reads that failed QC. We obtained transcript-level quantification using salmon 0.8.2^62^ (with --gcBias and --seqBias flags to account for potential biases) and the GRCh38 cDNA assembly release 87 downloaded from Ensembl. We converted transcript-level to gene-level count estimates, with estimates for 39037 genes based on Ensembl gene IDs. After QC, we retained expression estimates for 15994 genes with counts per million of 1 or higher in at least 10 samples. Limma-voom^63^ was used to remove heteroscedasticity from the estimated expression data. We tested genes for differential expression using limma in R (with lmFit and eBayes), based on a within-individual paired sample design. Significance was defined at 1% Benjamini-Hochberg FDR to correct for multiple testing. Of the 14408 genes with unique mapping of gene name and Ensembl ID, we took forward 1705 genes with significantly different abundance between intact and degraded cartilage at 1% FDR.

### Animal model data

The presence of abnormal skeletal phenotypes in mice was evaluated for all genes within 500kb of an osteoarthritis index variant and extracted from Open Targets^64^. This platform integrates all abnormal phenotype annotations for mutations in mouse genes reported in the literature and curated at MGI (URLs). Given the list of genes located less than 1 Mb away of the 64 genome-wide significant signals for osteoarthritis, abnormal skeletal system phenotypes from mutant mice were extracted systematically for all mouse orthologs of the human genes using the programmatic interface of the Open Targets platform (Supplementary Table 20). For instance, mutant mice homozygous for a targeted mutation of Smad3 (the ortholog of human SMAD family member 3) developed degenerative joint disease by progressive loss of articular cartilage^65^. Additional manual PubMed searches were conducted on selected genes to obtain information regarding animal models specific for osteoarthritis (Supplementary Table 19).

## DATA AVAILABILITY

All RNA sequencing data have been deposited to the European Genome/Phenome Archive (cohort 1: EGAD00001001331; cohort 2: EGAD00001003355; cohort 3: EGAD00001003354). Genotype data of the arcOGEN cases and UKHLS controls have been deposited at the European Genome-phenome Archive under study accession numbers EGAS00001001017 and EGAS00001001232, respectively.

## URLs

LDHub, http://ldsc.broadinstitute.org; OMIM, https://www.omim.org; Orphanet, http://www.orpha.net; HRC pre-imputation checking tool, http://www.well.ox.ac.uk/∼wrayner/tools/#Checking; MGI, http://www.informatics.jax.org; Open targets, https://www.opentargets.org; Understanding Society, https://www.understandingsociety.ac.uk; DEPICT, www.broadinstitute.org/depict; PASCAL www2.unil.ch/cbg/index.php?title=Pascal; DEPICT version 1 rel194 GitHub https://github.com/perslab/depict

## ACKNOWLEDGEMENTS

This research has been conducted using the UK Biobank Resource under Application Number 26041. This work was funded by the Wellcome Trust (206194). We are grateful to Roger Brooks, Andrew McCaskie, Jyoti Choudhary and Theodoros Roumeliotis for their contribution to the transcriptomic and proteomic data collection, and to Arthur Gilly for help with figures. The Human Research Tissue Bank is supported by the NIHR Cambridge Biomedical Research Centre. arcOGEN was funded by a special purpose grant from Arthritis Research UK (grant 18030). The UK Household Longitudinal Study was funded by grants from the Economic & Social Research Council (ES/H029745/1) and the Wellcome Trust (WT098051). UKHLS is led by the Institute for Social and Economic Research at the University of Essex. The survey was conducted by NatCen and the genome-wide scan data were analysed and deposited by the Wellcome Sanger Institute. Information on how to access the data can be found on the Understanding Society website https://www.understandingsociety.ac.uk. PICCOLO was developed by Karsten Sieber and Karl Guo. GDS and TRG receive funding from the UK Medical Research Council (MC_UU_00011/1 and MC_UU_00011/4). The authors would like to acknowledge Open Targets for enabling the collaboration on this work.

## AUTHOR CONTRIBUTIONS

UK Biobank association analyses: IT, LYA, RS, TJ, JH, EZ, JEG, KH, MK

arcOGEN analyses: arcOGEN, LS

Mendelian randomization: VH, JZ, RS, TG, GDS

Functional genomics: JMW, JEG, LMC, JS, LS, SB, DS, EZeggini

Translation work: LMC, JEG, NB, EZeggini

Manuscript writing: IT, KH, LS, JEG, LMC, RS, EZeggini

## COMPETING INTERESTS

IT, JEG, TJ, LYA, JH, NB, RS, LMC are employees of GlaxoSmithKline and may own company stock. TRG receives research funding from GlaxoSmithKline and Biogen. VH is funded by a research grant from GlaxoSmithKline.

